# Exotic bifurcations in three connected populations with Allee effect

**DOI:** 10.1101/2021.02.03.429609

**Authors:** Gergely Röst, AmirHosein Sadeghimanesh

**Author notes:** http://www.math.u-szeged.hu/~rost/ (Gergely Röst).

## Abstract

We consider three connected populations with strong Allee effect, and give a complete classification of the steady state structure of the system with respect to the Allee threshold and the dispersal rate, describing the bifurcations at each critical point where the number of steady states change. One may expect that by increasing the dispersal rate between the patches, the system would become more well-mixed hence simpler. However, we show that it is not always the case, and the number of steady states may (temporarily) increase by increasing the dispersal rate. Besides sequences of pitchfork and saddle-node bifurcations, we find triple-transcritical bifurcations and also a sun-ray shaped bifurcation where twelve steady states meet at a single point then disappear. The major tool of our investigations is a novel algorithm that decomposes the parameter space with respect to the number of steady states and find the bifurcation values using cylindrical algebraic decomposition with respect to the discriminant variety of the polynomial system.

## 1 Introduction

In population dynamics, Allee effect refers to a situation when a population has lower growth rate at small densities. There are several biological mechanisms that can create this effect, and its significance in ecology has been widely demonstrated [2].

The simplest mathematical example of an Allee effect is given by the cubic growth model

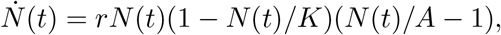

where *N* (*t*) is the population size, 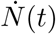 is its time derivative, *r* is the intrinsic growth rate, *K* is the carrying capacity, and *A* ∈ (0, *K*) is called the Allee threshold. From a dynamical systems point of view, this is also one of the simplest systems with bistability: populations above the Allee threshold persist and converge to the carrying capacity, whereas populations that fall under the Allee threshold go extinct. With the transformation 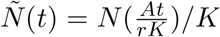, after dropping the tilde and using the notation *b* = *A/K*, the cubic growth equation can be rescaled to

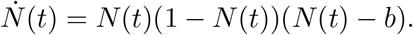

While this equation is indeed simple, things get interesting when we consider the spatial distribution of a species exhibiting Allee effect and living at different habitats which are connected by spatial dispersal. In that case, a large number of steady states emerge with a variety of configurations depending on the network of connections between the patches [11]. Multiple attractors and long transients have been identified in discrete time models for spatially structured populations with Allee effect ([5], [9]). When there are only two connected patches, up to nine steady states can coexist, and the bifurcations of this planar system have been investigated for a continuous time model in [7]. The goal of our work is to provide a complete description of the bifurcations for populations dispersed at three connected patches.

To achieve this goal we first translate the question of interest, namely for what choice of parameters the number of steady states changes, to an algebraic question. Then we adopt a symbolic method called Cylindrical Algebraic Decomposition (CAD) to decompose the parameter space with respect to the discriminant variety of the system. The discriminant variety of a parametric polynomial system of equations is the Zariski closure of the locus of the parameter points for which the system has a solution with multiplicity greater than one. This method is implemented in a Maple package `RootFinding[Parametric]`. But the computation for the model in this paper is not feasible on our computer. Therefore we introduce a new algorithm, Algorithm 1, which is combining a numeric search algorithm with the above symbolic tool.

In recent years, various combinations of symbolic and numeric algorithms to study steady states of biological networks have been developed. In [1, 4], CAD and Real Triangularization have been used to count the number of steady states for parameter values coming from a grid of sample points, giving an approximate identification of the decomposition of the parameter space with respect to the number of steady states of the system. In that approach, the discriminant variety is not used. On the other hand, [8] uses numerical homotopy methods to decompose the parameter space with respect to the discriminant variety. Our approach is different from both mentioned works, since we use the discriminant variety, but not numerical homotopy methods. Instead we adopt a bisection search for finding minimum and maximum of some distance functions being defined in section 4.

Our paper illustrates how such methods and tools can be used to give a comprehensive description of the bifurcations in biological models formulated as polynomial ODE systems, such as population dynamics with spatial dispersal.

## 2 Three patch model

In the following sections we consider a compartmental model consisting of three patches. Depending on the context a patch can be interpreted as a habitat, a cellular compartment, a well in a microplate etc. For *i* = 1, 2, 3 let *N*_*i*_(*t*) denote the population at the *i*th patch at time *t*. We drop the emphasis on *t* and simply write *N*_*i*_. The evolution of the populations in the three patches is modeled by the following ODEs in 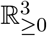.

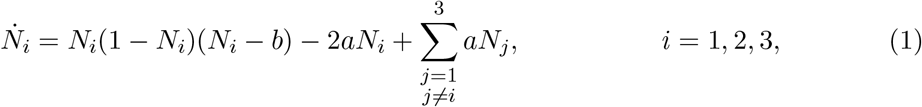

where 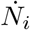 stands for 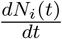. The parameters *b* ≥ 0 and *a* ≥ 0 correspond to the Allee threshold and the dispersal rate between patches respectively. In this model all three patches are identical and connected reversibly with equal dispersal rates. In another word, the connection graph of the patches is a complete simple labeled digraph with the same labels.

The *steady states* of the ODE system (1) for fixed values of the parameters *a* and *b* are the non-negative solutions to the polynomial system of equations obtained by letting 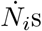 equal to zero. The goal of the next section is to study the number of steady states of this three patches model. Speaking algebraically, we consider the following parametric polynomial system of equations in three variables *N*_1_, *N*_2_, *N*_3_ and two parameters *a, b*.

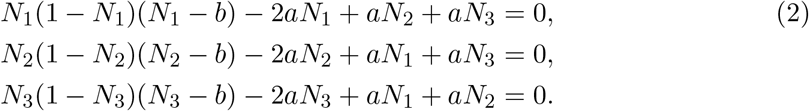

Then we want to determine what are the possible number of non-negative solutions to (2) and describe the conditions on the parameters for each case. Before we proceed to the next section, we prove two lemmas.

### Lemma 2.1

*Consider the following ODE system in n variables N*_*i*_, *i* = 1, *…, n and two parameters a, b*.

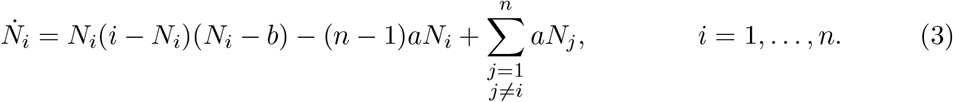

*The number of steady states of* (3) *for the parameter point* 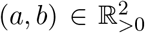 *is equal to the number of steady states for the parameter point* 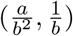.

*Proof*. Using the change of variables 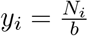, *i* = 1, *…, n*, we have

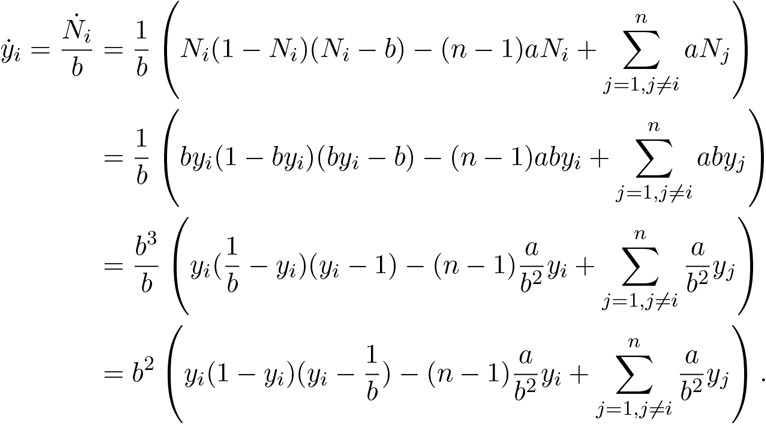

Hence for any solution (*x*_1_, *…, x*_*n*_) of the system for the parameter point (*a, b*), we have a solution 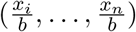 for the parameter point 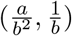 and vice versa. This proves that the number of steady states (equilibria) of the system is the same for the two choices of the parameter points.□

### Lemma 2.2

*Consider the ODE system in n variables N*_*i*_, *i* = 1, *…, n and two parameters a, b given in* (3). *The number of steady states of the system for the parameter point* (*a, b*) ∈ ℝ_*>*0_×(0, 1) *is equal to the number of steady states for the parameter point* (*a*, 1−*b*).

*Proof*. Using the change of variables *y*_*i*_ = 1 − *N*_*i*_, *i* = 1, *…, n*, we have

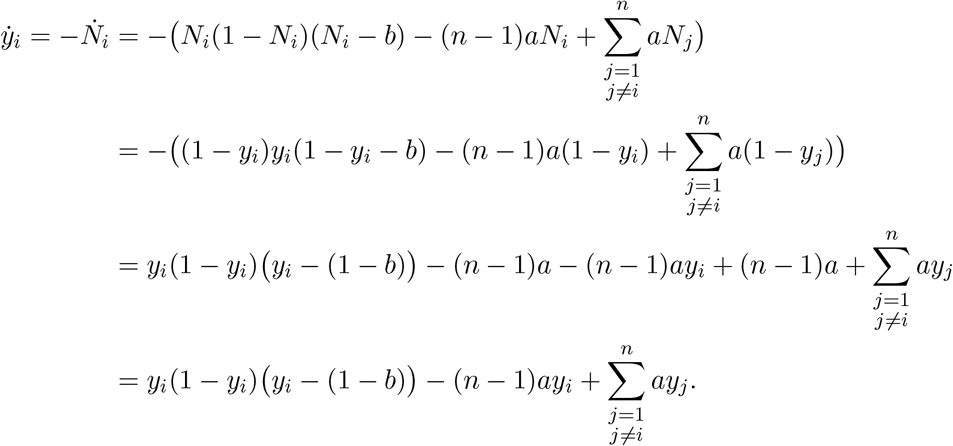

Hence for any solution (*x*_1_, *…, x*_*n*_) of the system for the parameter point (*a, b*), we have a solution (1 − *x*_1_, *…*, 1 − *x*_*n*_) for the parameter point (*a*, 1 − *b*) and vice versa. Since for *b <* 1 we have *x*_*i*_ ≤ 1, this proves that the number of steady states of the system is the same for the two choices of the parameter points. □

By Lemmas 2.1 and 2.2, it is enough to study the number of steady states of (3) only on the parameter region 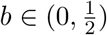.

## 3 Algebraic Preliminaries

Here we introduce the necessary algebraic concepts that we later use to study the behavior of the polynomial ODE system in (1).

Let *R* = ℚ [*k*_1_, *…, k*_*m*_][*x*_1_, *…, x*_*n*_] be the ring of parametric polynomials in *n* variables, *x*_1_, *…, x*_*n*_, and *m* parameters, *k*_1_, *…, k*_*m*_, with rational coefficients. For a set *F* ⊆ *R* and a choice of parameter values (*k*_1_, *…, k*_*m*_) ∈ ℝ^*m*^, let *V*_*k*_(*F*) denote the real solution set to the system of equations {*f* = 0 | *f* ∈ *F*}. The discriminant variety associated with *F* is the Zariski closure of the set of points in the parameter space for which the system *V*_*k*_(*F*) has a solution with multiplicity higher than one. Hereafter, assume that *F* = {*f*_1_, *…, f*_*n*_} and the ideal generated by *F* is a zero dimensional ideal. In this case the discriminant variety is the solution set to any basis of the following ideal.

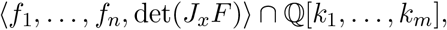

where *J*_*x*_*F* is the Jacobian matrix of *F* with respect to the variables and *f*_*i*_s are considered as polynomials in ℚ[*k*_1_, *…, k*_*m*_, *x*_1_, *…, x*_*n*_]. Denote the discriminant variety by *V*_*D*_.

In addition to the equations, assume that there are inequality constraints *g*_*i*_ ≺_*i*_ 0 where *G* = {*g*_1_, *…, g*_*s*_} ⊂ *R* and ≺_*i*_ can be any of the relations ≠, *<, >*, ≤ or ≥. For any choice of parameters (*k*_1_, *…, k*_*m*_) ∈ ℝ ^*m*^, let *V*_*k*_(*F, G*) be the set of points in *V*_*k*_(*F*) that satisfy the inequalities *g*_*i*_ ≺ 0. By changing the values of the parameters on a continuous path, the number of points in *V*_*k*_(*F, G*) can not change and these points move continuously unless at least one of the following events occur.

i. Two points in *V*_*k*_(*F*) meet and become a pair of non-real complex conjugates, therefore leaving *V*_*k*_(*F*). Or the opposite, two complex conjugate solutions of *f*_1_ = … = *f*_*n*_ = 0 meet and become two real solutions, therefore entering *V*_*k*_(*F*).
ii. A point in *V*_*k*_ (*F*) crosses a hypersurface defined by one of the inequality constraints, g_*i*_ = 0, and leave or enter *V*_*k*_ (*F, G*).
iii. A point from *V*_*k*_(*F*) leaves the affine space, i.e. becomes a solution at infinity.

Since event (i) implies having a solution with multiplicity at least two, this means the parameter path must cross *V*_*D*_. Note that the opposite is not necessarily true. Denote the Zariski closure of the two sets of the parameter points for which the events (ii) and (iii) happen with *V*_≺_ and *V*_∞_ respectively. It is easy to see that *V*_≺_ is subset of the solution set to any basis of the following ideal.

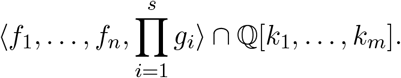

There is more care needed in order to compute *V*_≺_ than just computing the above ideal. We refer the reader to [12]

Let *x*_0_ be a new auxiliary variable. For a polynomial *f* ∈ *R* of the total degree deg(*f*) with respect to only *x*_1_, *…, x*_*n*_, define *f*^*h*^ to be the polynomial that each of its terms, say *t*, is a same term as in *f* multiplied by 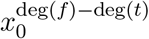. The polynomial *f*^*h*^ is called the homogenization of *f* and is an element in the ring ℚ [*k*_1_, *…, k*_*m*_][*x*_0_, *x*_1_, *…, x*_*n*_]. In addition, we have that 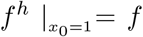. One can see that *V*_∞_ is subset of the union of the solution sets to any basis of the following ideals.

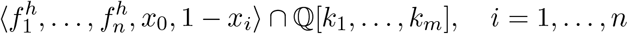

Similar to *V*_≺_, we refer the reader to [12] for how to compute *V*_∞_.

It is common to modify definition of discriminant variety to be *V*_*D*_ ∪ *V*_≺_ ∪ *V*_∞_ and call the points on the discriminant variety, *critical points*. The key property of the discriminant variety is that the number of points in *V*_*k*_(*F, G*) is invariant on the connected components of the complement of this variety.

Using discriminant variety we took a system in *n* + *m* indeterminates and make a new set of polynomials in *m* indeteminates. What is left is to decompose the parameter region with respect to this variety. To this end a common tool is *Cylindrical Algebraic Decomposition* (CAD). The output of CAD algorithm for the discriminant variety is a collection of closed and open regions called *cells*, each of them entirely inside the discriminant variety or a connected component of the complement of the discriminant variety. The number of elements in *V*_*k*_(*F, G*) is invariant on each cell. Another important key fact about CAD is that the number of cells in the output is finite. Thus it is enough to pick up one sample point from each cell and solve the system for the parameter values determined by the sample point. To read more about this method see [12, 13]. This method is implemented in a Maple package called `RootFinding[Parametric]` [6]. This package computes the open CAD, that is the output consists of only the open cells. In this paper, we simply say “CAD” instead of “open CAD of the parameter space with respect to the disciminant variety”.

## 4 Multiple steady states via the CAD method

As illustrated in Section 2, the goal of this section can be simplified to the study of the number of solutions to the system of equations (2) together with the inequalities given below:

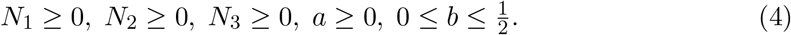

Section 3 indicates that one approach to achieve our goal is to use CAD. Unfortunately Maple’s `RootFinding[Parametric]` package can not compute the CAD for the system made by equations (2) and (4) on our computer^1^ with both *a* and *b* as free parameters. Attempting to compute *V*_*D*_ using Gröbner basis computation with Singular [3] was not feasible with our computational resources.

Thus we have to find an alternative approach to overcome the complexity of CAD for our model with two free parameters. Instead of letting both parameters *a* and *b* to be free, fix the value of *b* to be 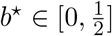. The 1-dimensional CAD with respect to *a* consists of a finite number of disjoint open intervals. The union of these open intervals is equal to R_*>*0_ minus a finite number of points. A pair (*a*^⋆^, *b*^⋆^) where *a*^⋆^ is one of such points, is on the intersection of the line *b* = *b*^⋆^ with the 2-dimensional discriminant variety. Hence by computing 1-dimensional CADs of the model for different values of *b*, one can construct an approximation of the 2-dimensional CAD.

Following this idea, we chose 101 equally distanced values for *b* between and including 0 and 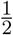. Then we compute the 1-dimensional CAD with respect to *a* for each of the choices of *b*. For each of these values of *b* we get five critical values for *a*. Denote these five values by *α*_*i*_, *i* = 1, *…*, 5, ordered from the smallest number to the largest and note that the value of *α*_*i*_ is dependent on the choice of *b*. Connecting *α*_*i*_s we get five curves as shown in Figure 1.

**Figure 1.**
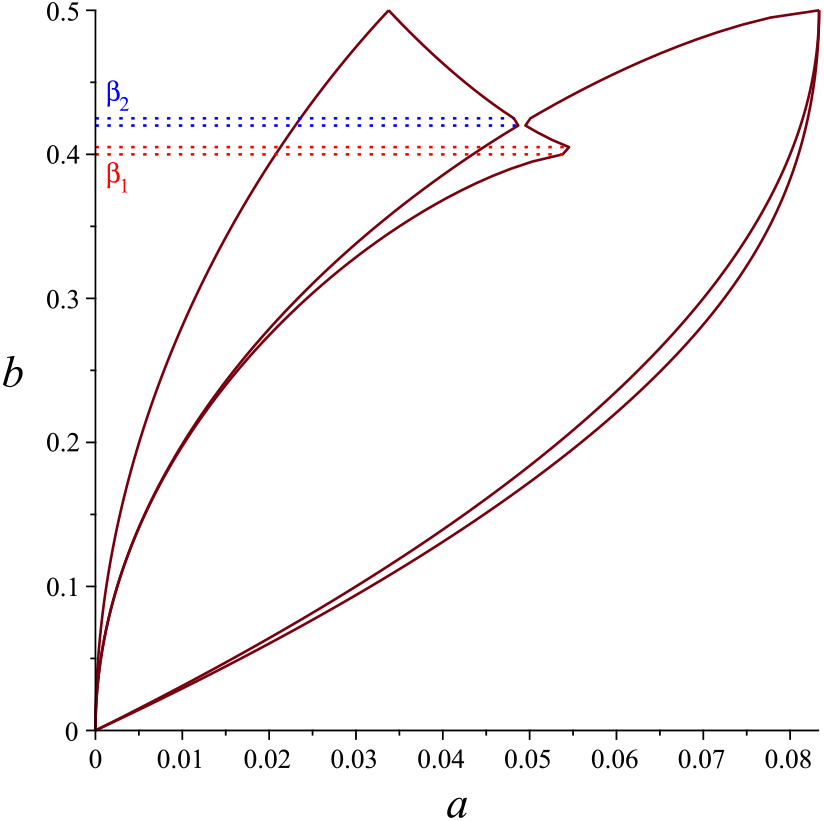
For 101 equally distanced values of *b* between and including 0 and 0.5, the 1-dimensional CAD of the system made by (2) and (4) after substituting the value of *b* gives five closed cells, the singletons {*α*_*i*_} and six open cells, the intervals made by removing *α*_*i*_s from [0, ∞). Connecting *α*_*i*_s for different *b*’s gives the five curves depicted in this figure. The figure suggests that these curves have intersection or sharp angle at four values of *b*; 0, 1*/*2 and two values in between denoted by *β*_1_ and *β*_2_.

Figure 1 suggests that there are four possible values for *b*; 0, *β*_1_, *β*_2_ and 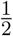 where the number of critical values of *a* in the 1-dimensional CADs may change. We refer to these values as critical values for *b*.

- At *b* = 0 we have that *α*_1_ = … = *α*_5_ = 0.
- At 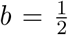 we have that 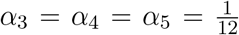 and *α*_1_ = *α*_2_ = *c* where *c* is the only positive real root of 27648*x*^4^ − 576*x*^2^ + 48*x* − 1 = 0 which with seven digits after the decimal point is 0.0337670.

The two other points are guessed to have the following properties.

- At b = β _1_, the distance of α _2_ and α _3_ becomes maximum on (0, β _2_)
- At b = β _2_, α _2_ = α _3_.

To investigate the behavior of the system around *b* = *β*_1_ and *b* = *β*_2_ we introduce a new algorithm.

### 4.1 Searching for the critical values of *b*

Define Δ(*b*) to be *α*_3_ − *α*_2_ for a given *b*. Then Δ seems to be strictly increasing before *β*_1_ and strictly decreasing after *β*_1_ and therefore it can not have any local maximum other than *β*_1_ on (0, *β*_2_). From Figure 1, *β*_1_ should belong to (4, 4.05). To find *β*_1_ with a desired accuracy we adopt a bisection search strategy.

Let *b*_1_ = 4 and *b*_2_ = 4.05 and *b*_3_ = (*b*_1_ + *b*_2_)*/*2. If Δ(*b*_3_) *<* Δ(*b*_2_), then *β*_1_ can not belong to (*b*_1_, *b*_2_) otherwise we will have another local maximum in (*b*_3_, *b*_2_). Therefore we can restrict our search domain from (*b*_1_, *b*_2_) to (*b*_2_, *b*_3_). Similarly if Δ(*b*_3_) *>* Δ(*b*_1_), then we can restrict our search domain to (*b*_1_, *b*_3_).

Clearly if none of these two happen, the only possibility is to have Δ(*b*_3_) being greater than both of Δ(*b*_1_) and Δ(*b*_2_). In this case we define two new midpoints; *b*_4_ = (*b*_1_ + *b*_3_)*/*2 and *b*_5_ = (*b*_3_ + *b*_2_)*/*2. If Δ(*b*_4_) *>* Δ(*b*_3_), then *β*_1_ can not belong to (*b*_3_, *b*_2_), otherwise we have another local maximum in (*b*_1_, *b*_3_). Therefore we can restrict our search domain to (*b*_1_, *b*_3_). Similarly if Δ(*b*_5_) *>* Δ(*b*_3_), then we can restrict the search domain to (*b*_2_, *b*_3_).

If neither of these two happen, then we have Δ(*b*_4_) *<* Δ(*b*_3_) and Δ(*b*_5_) *<* Δ(*b*_3_). In this case *β*_1_ can not belong to (*b*_1_, *b*_4_) ∪ (*b*_5_, *b*_2_), otherwise we have another local maximum in (*b*_4_, *b*_5_). Therefore we can restrict the search domain to (*b*_4_, *b*_5_).

Hence in each five possible cases, it is possible to replace (*b*_1_, *b*_2_) with a sub-interval of a half length. This proves that by continuing this process we are guaranteed to reach an interval of length smaller than a given precision value and hence finding *β*_1_ with a desired accuracy. Algorithm 1 summarises this bisecting search strategy. At *b* = *β*_2_ we have that *α*_2_ = *α*_3_. In other words Δ(*b*) = 0 which is a minimum on (*β*_1_, 0.5). Since Δ(*b*) seems to be strictly decreasing before *β*_2_ and strictly increasing after *β*_2_, one can use the same algorithm as for finding *β*_1_, but reversing the inequality side in the conditions and letting *b*_1_ = 4.2 and *b*_2_ = 4.25. We implemented Algorithm 1 on Maple and found *β*_1_ = 0.4014757 and *β*_2_ = 0.4215113 with seven digits of accuracy after the decimal point. The computation took 879 and 1591 seconds for finding *β*_1_ and *β*_2_ respectively.

Computing the 1-dimensional CADs for the *b* values arising in the steps of Algorithm 1 for *β*_1_ and *β*_2_ gives Figures 2. The plot in Figure 2a shows that in a small neighborhood before *β*_1_ there is a small region missed by Figure 1. It also shows that there is a fifth critical value for *b* near to *β*_1_. Denote this value with 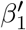. For the values of *b* between *β*_1_ and 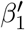, instead of 5 critical values for *a*, there are 7 critical values. Ordering the critical values of *a* from the smallest to the largest, the two new values are the 4th and the 5th which we show them by 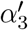 and 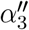 and use the same notation *α*_*i*_, *i* = 1, *…*, 5 for the rest of them. To find 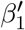 with desired accuracy, we again use Algorithm 1, but with a different definition of Δ. Let Δ(*b*) = *α*_3_ − *α*_2_ if the 1-dimensional CAD has 5 critical values for *a*, otherwise let 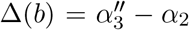. By Figure 2a, 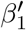 should belong to (0.40125, 0.40140625). The value of 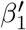 with seven digits after the decimal point is 0.4013889.

**Figure 2.**
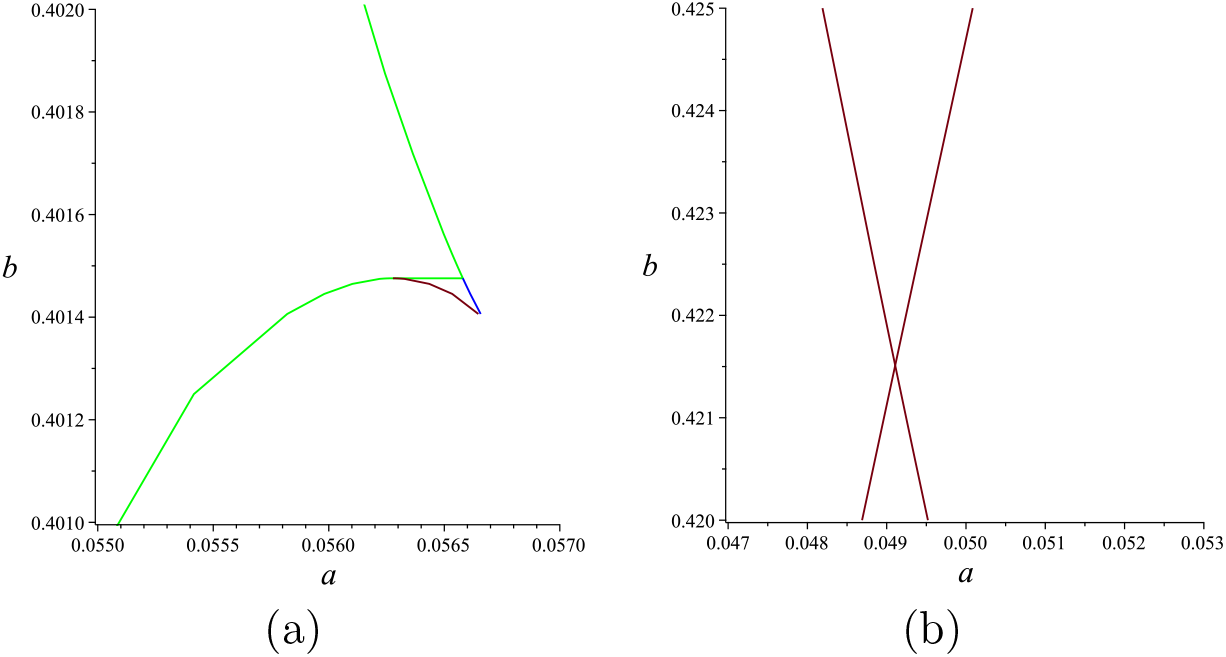
(a) For the values of *b* arising in the steps of Algorithm 1 for finding *β*_1_, the 1-dimensional CAD of the system made by (2) and (4) is computed. For some values in a small interval instead of 5 critical values for *a*, we get 7 critical values. The two new critical values are very close to *α*_3_. Denote the two new critical values by 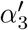 and 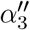. The green, red and blue curves are made by connecting *α*_3_, 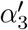 and 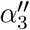 respectively. (b) For the values of *b* arising in the steps of Algorithm 1 for finding *β*_2_, the 1-dimensional CAD of the system made by (2) and (4) is computed. The two crossing curves are made by connecting values of *α*_2_ and *α*_3_.

#### Algorithm 1

The bisecting search algorithm for finding *β*_1_ (and 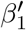) with the precision *E*. To find *β*_2_, direction of the inequalities in the if statements of steps 3, 4, 6 and 8 should be reversed.

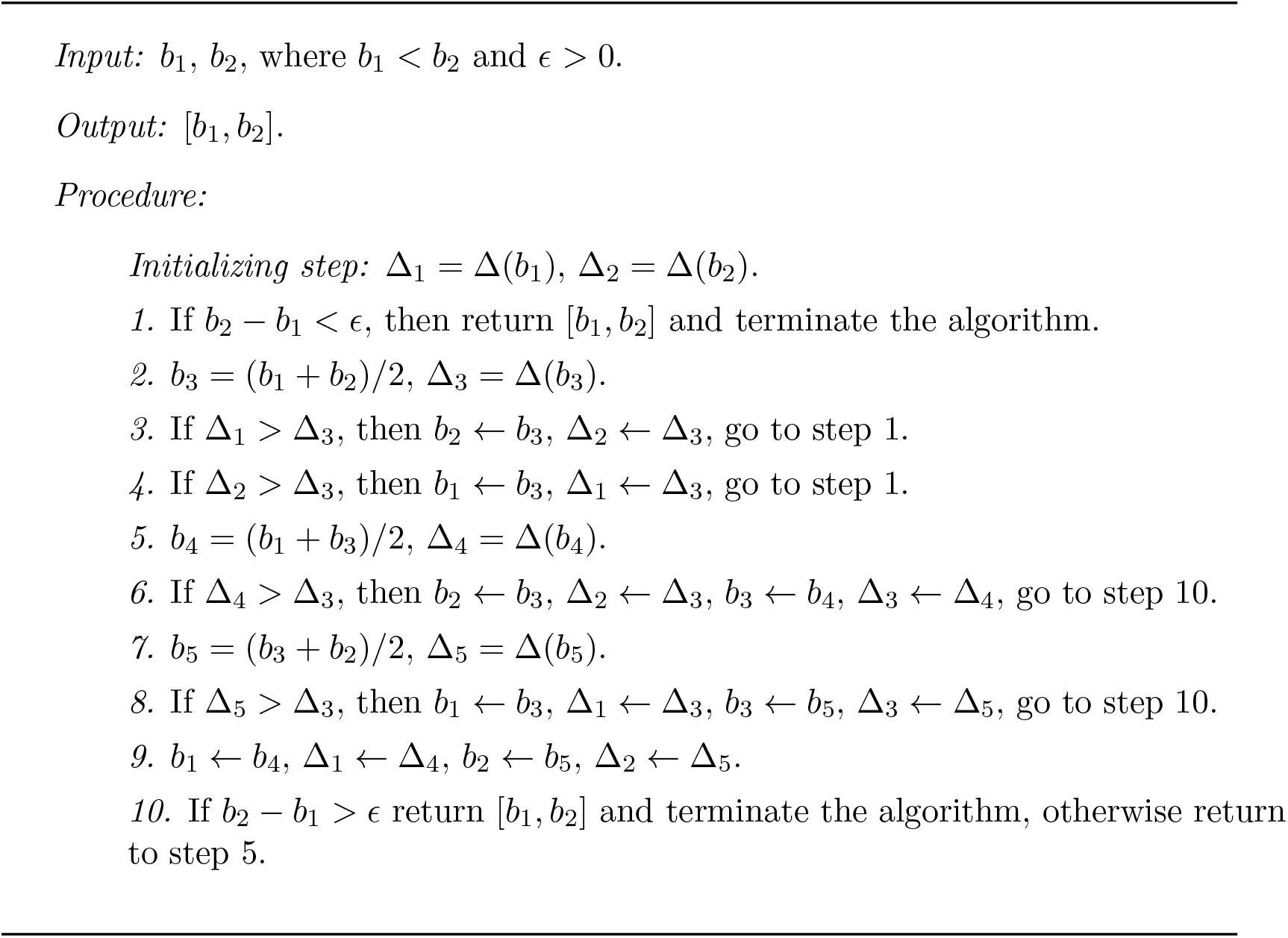

### 4.2 The two dimensional CAD of the model

We found 5 critical values for *b*; 0, 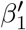, *β*_1_, *β*_2_ and 1*/*2. The behavior of the system differ depending on the value of *b* being equal to these critical values or between them. There are 7 possible sequences of the number of steady states for a choice of *b* by increasing value of *a*. Table 1 shows a complete description of the regions in each sequence. Figure 3 shows a plot of regions colored with respect to the number of steady states. A brief description is listed below.

**Table 1:**
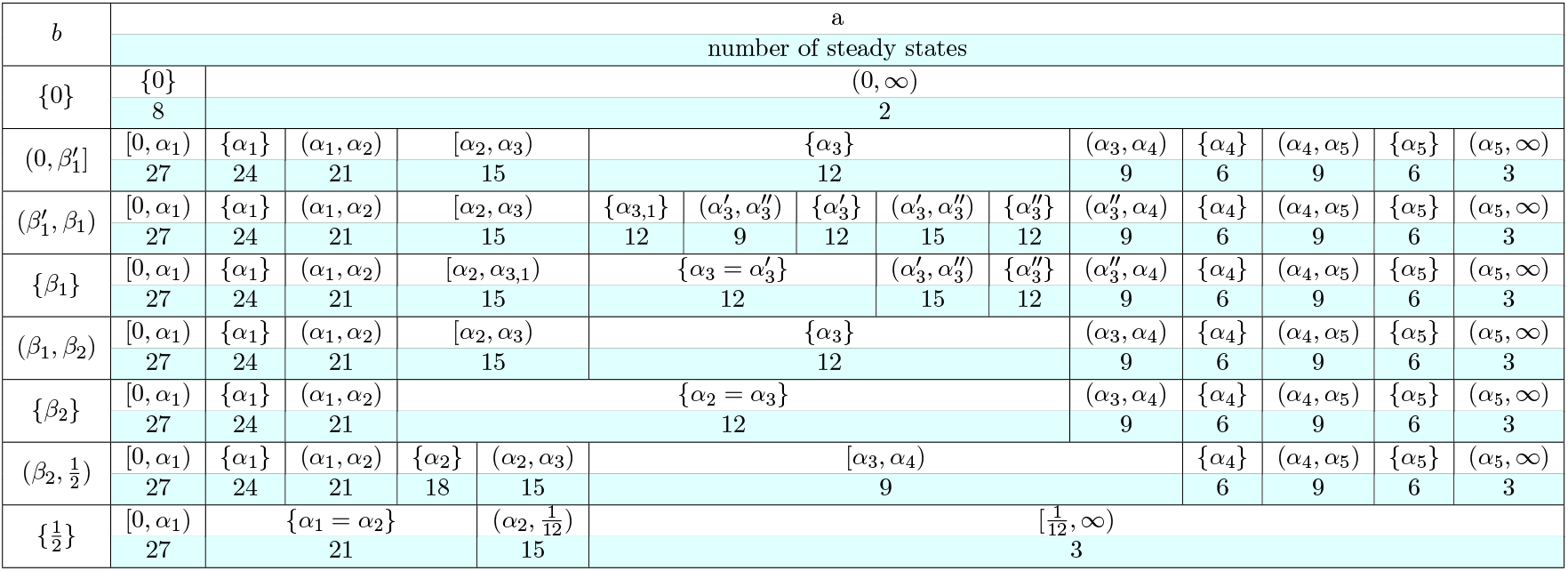
Number of non-negative steady states of the system (1) for the parameters *a* ≥ 0, 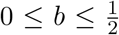. The first column shows the interval of *b*. Each row for the intervals of *b* has two sub-rows, the first sub-row shows the interval of *a* and the second sub-row states the number of steady states.

**Figure 3.**
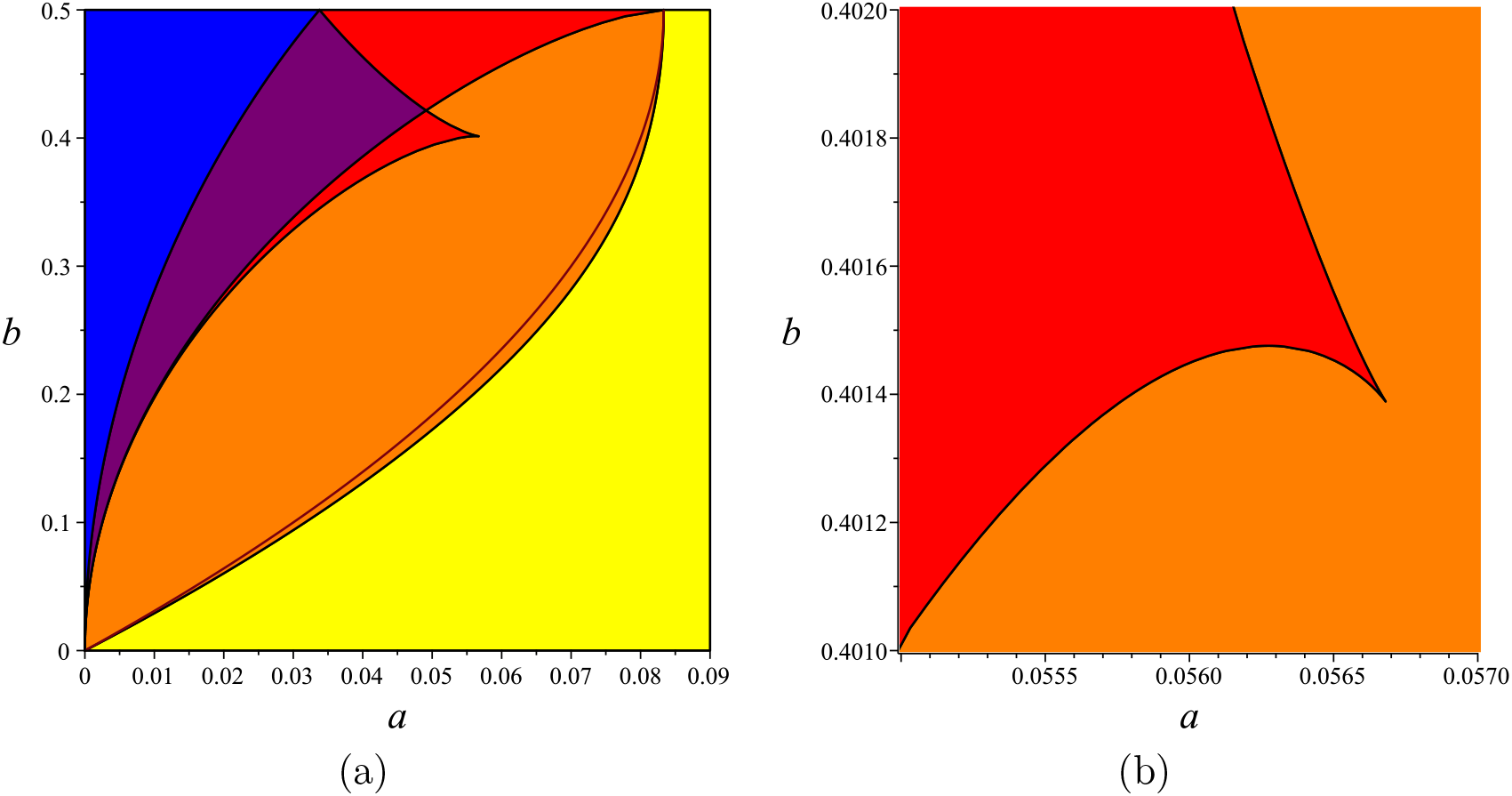
(a) Partitioning the parameter region 0 ≤ *a*, 0 ≤ *b* ≤ 0.5 for the system of equations (2) with respect to the number of non-negative solutions. For *a* = 0 the system has 27 solutions. On the blue, purple, red, orange, yellow colored regions the system has 27, 21, 15, 9 and 3 solutions respectively. On the red curve inside the orange colored region, the system has 6 solutions. For the number of solutions on the boundaries of the regions see Table 1. (b) Enlargement of the region 0.055 *< a <* 0.057, 0.401 *< b <* 0.402.

1. *b* = 0; All five critical values of *a* become equal *α*_1_ = … = *α*_5_ = 0. Possible number of steady states are 2, 8.
2. 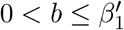, *β*_1_ *< b < β*_2_; All *α*_*i*_’s are separate. Possible number of steady states are 3, 6, 9, 12, 15, 21, 24, 27.
3. 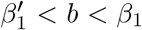; There are two more critical values for *a*; 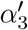 and 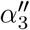. Possible number of steady states are 3, 6, 9, 12, 15, 21, 24, 27.
4. *b* = *β*_1_; There is one more critical value for *a*; 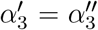. Possible number of steady states are 3, 6, 9, 12, 15, 21, 24, 27.
5. *b* = *β*_2_; The second and third critical values of *a* collide, *α*_2_ = *α*_3_. The possible number of steady states are 3, 6, 9, 12,21,24,27.
6. 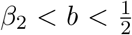; All *α*_*i*_’s are separate. Possible number of steady states are 3, 6, 9, 15, 18, 21, 24, 27.
7. 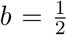; The first and second critical values collide, as well as the third, fourth and fifth, *α*_1_ = *α*_2_ and 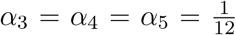.Possible number of steady states are 3, 15, 21, 27.

## 5 Bifurcation sequences

There are five possible types of bifurcation events for the system (1). Four of these bifurcation events are depicted in Figure 4. Figures 6 and 7 shows the behavior of the steady states for fixed *b* and increasing *a*. Stability of the steady states are shown in the plots on the right side, a steady state is colored red or pink if it is stable or unstable respectively. The steady states in the plots on the left side are colored with respect to the value of *a* which can be read from the color bar next to it. To describe the behavior of the system we consider several cases listed below.

**Figure 4.**
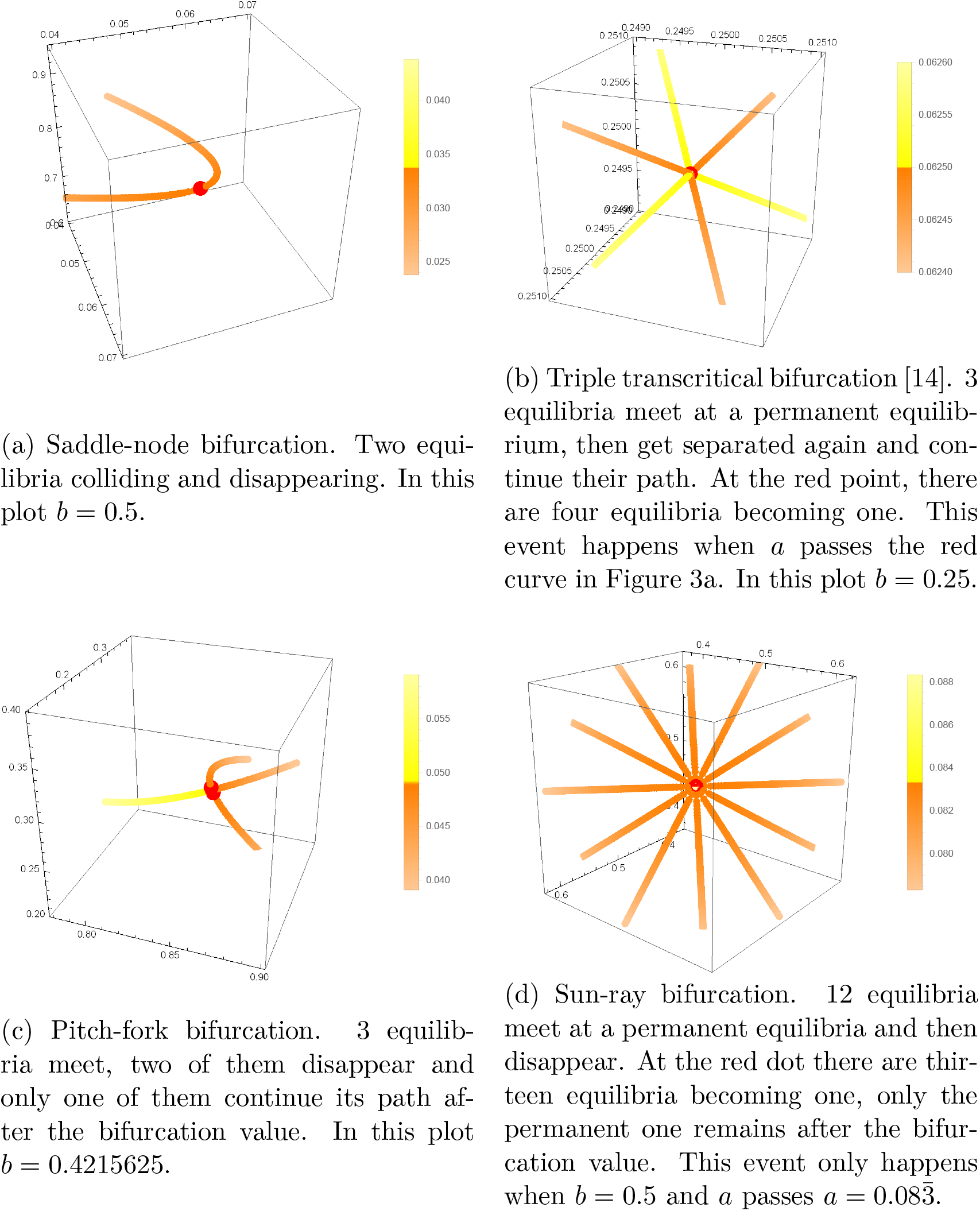
Several examples of the possible bifurcations of the system (1) are depicted. Orange, yellow and red colors are used to show the steady states before, after and at the bifurcation value of *a* respectively. The color bars in the right side of the plots show the interval for the values of *a*

**Figure 5.**
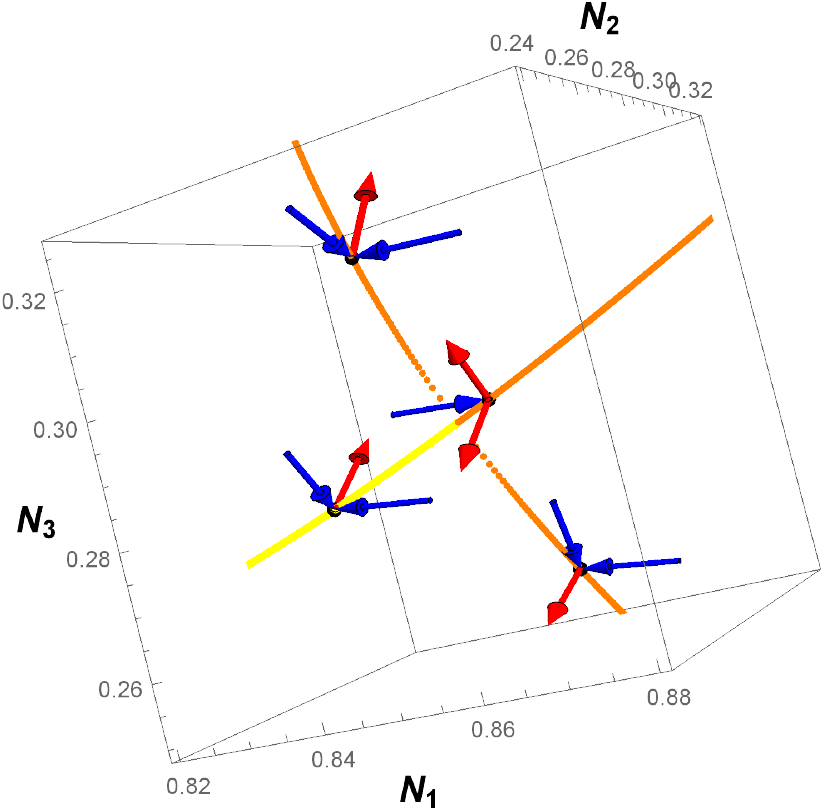
The pitchfork bifurcation at Figure 4c. The Allee effect parameter is fixed to 0.4215625 and the strength of the connectivity between parameters is varying from 0.0391 to 0.0591. The bifurcation occurs at *a* 0.04912148135. Steady states for the values of *a* before and after the bifurcation value are colored by orange and yellow respectively. For one choice of *a* before the bifurcation value, the eigenvectors of Jacobian matrix of the system at the three steady states are depicted as vectors. The direction of the vectors are toward the steady state if the corresponding eigenvalue has a negative real part and the opposite direction otherwise. Stabilitity also can be read from the colors of these vectors other than their directions. Blue is used for the stable directions and red for unstable ones. The same is done for one value of *a* after the bifurcation value. All four points have one similar stable and one unstable directions and the difference is only along the third direction. Before the bifurcation event the third direction is stable for two steady states and unstable for the middle steady state, after the bifurcation the third direction is stable.

**Figure 6.**
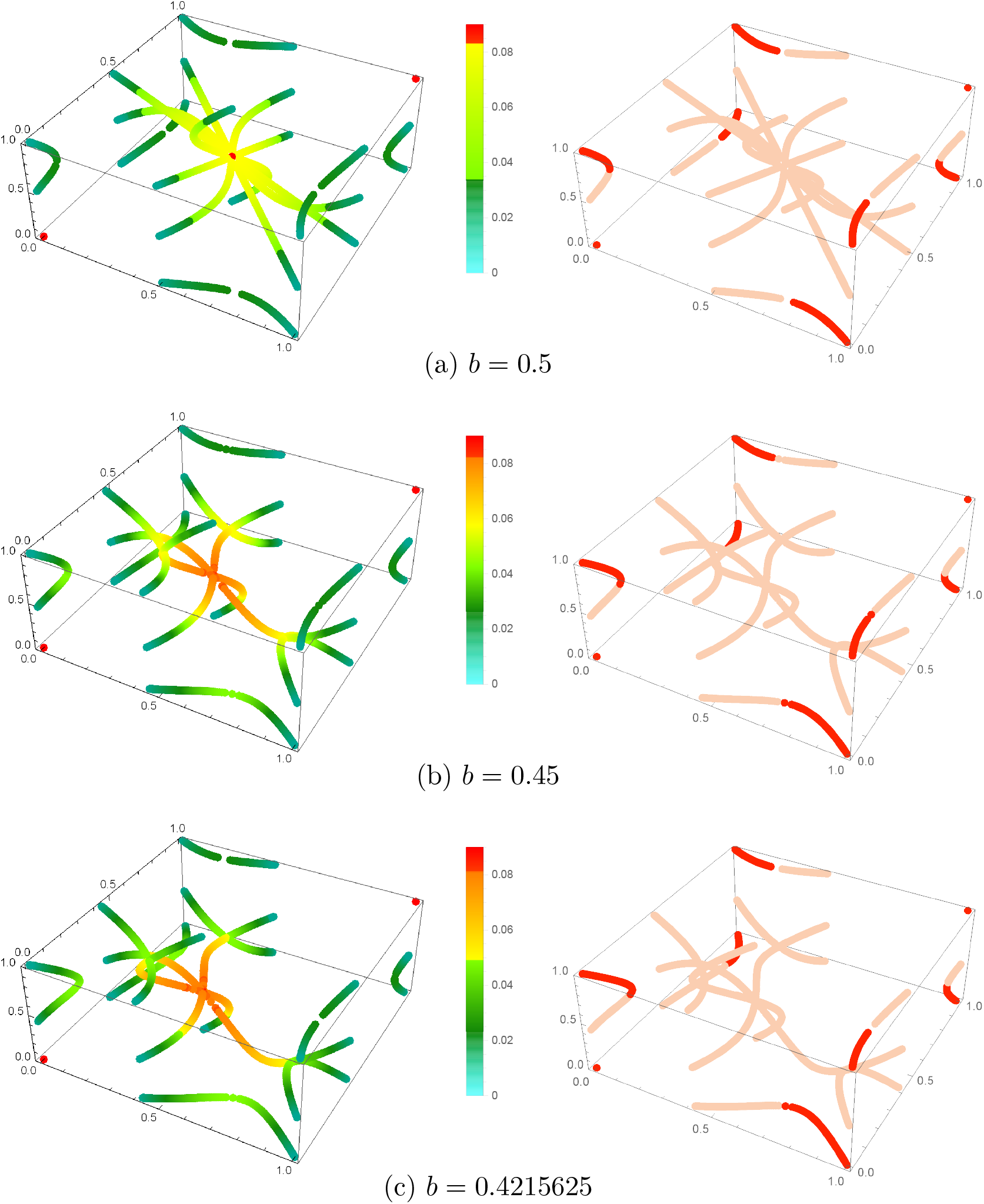
Sequences of the steady states for fixed values of *b*, but varying values of *a*. In the left side the steady state points are colored with respect to the value of *a* that can be read from the color bar next to the plots. In the right side the steady states are colored with respect to their stability. Stable steady states are colored by red, whereas the unstable ones are colored by pink. Continued at Figure 7.

**Figure 7.**
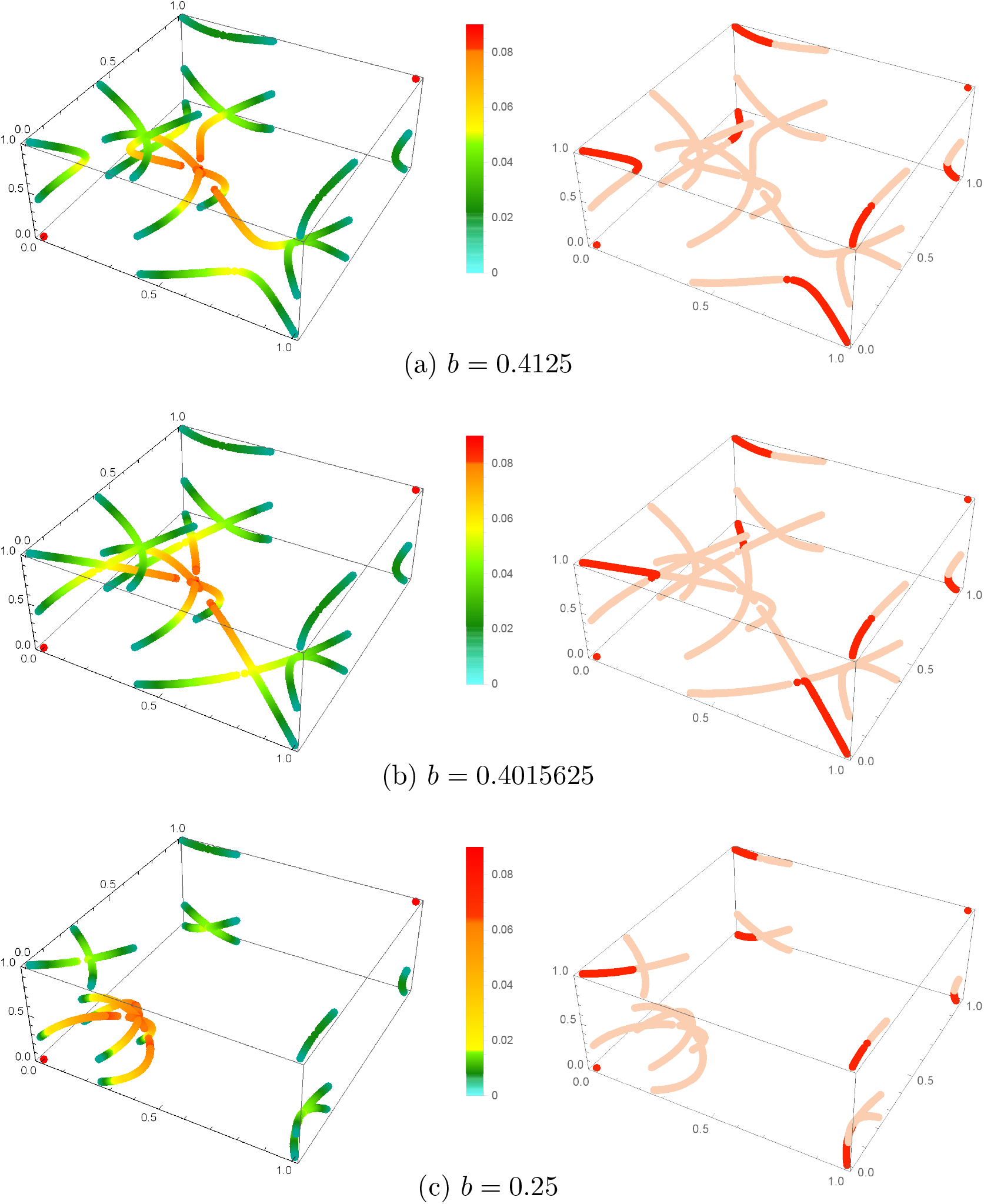
Continuation of Figure 6. Sequences of the steady states for fixed values of *b*, but varying values of *a*. In the left side the steady state points are colored with respect to the value of *a* that can be read from the color bar next to the plots. In the right side the steady states are colored with respect to their stability. Stable steady states are colored by red, whereas the unstable ones are colored by pink.

1. 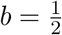; In this case there are two bifurcation points.
  a. 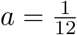; Six pairs of steady states collide and disappear in saddle-node bifurcation events. In each pair a steady state is stable and the other is unstable.
  b. *a* = *α*_3_ = *α*_4_ = *α*_5_; 12 unstable steady states move toward the middle stable steady state and after reaching it, all except the stable steady state disappear.
2. 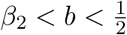; In this case there are five bifurcation points.
  a. *a* = *α*_1_; Three saddle-node events happen.
  b. *a* = *α*_2_; Another three saddle-node events happen.
  c. *a* = *α*_3_; Three pitch-fork events happen. In each pitch-fork, three unstable steady states collide, then two disappear and only one continue its path.
  d. *a* = *α*_4_; Three unstable steady states move toward the middle stable steady state, reach this point and then pass from it and continue their paths. It is a triple transcritical bifurcation event.
  e. *α*_5_; Three saddle-node events happen. At each saddle-node event, both steady states are unstable.
3. *b* = *β*_2_; In this case there are four bifurcation points.
  a. *a* = *α*_1_; Three saddle-node events happen.
  b. *a* = *α*_2_ = *α*_3_; Three saddle-node events and three pitch-fork events happen at the same time.
  c. *a* = *α*_4_; The triple transritical event happens.
  d. *a* = *α*_5_; Three saddle-node events happen.
4. *β*_1_ *< b < β*_2_; In this case there are five bifurcation points.
  a. *a* = *α*_1_; Three saddle-node events happen.
  b. *a = α* _2_; *Three pitch-fork events happen*.
  c. *a = α* _3_; *Three saddle-node events happen*.
  d. *a = α* _4_; *The triple transcritical event happen*.
  e. *a = α* _5_; Three saddle-node events happen.
5. *b* = *β*_1_; In this case there are six bifurcation points.
  a. *a* = *α*_1_; Three saddle-nodes events happen.
  b. a = α _2_; Three pitch-fork events happen.
  c. 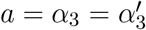; Three pairs of a stable and an unstable steady states collide and then continue their paths.
  d. 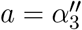; Three saddle-node events happen.
  e. a = α _4_; The triple transcritical event happen.
  f. a = α _5_; Three saddle-node events happen.
6. 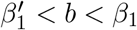; In this case there are seven bifurcation points.
  a. *a* = *α*_1_; Three saddle-nodes events happen.
  b. a = α _2_; Three pitch-fork events happen.
  c. a = α _3_; Three pairs of a stable and an unstable steady states collide and become a pair of complex conjugate points. So three saddle-node events happen.
  d. 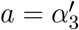; The three pairs of complex conjugate points of the previous step meet again and become three pairs of steady states.
  e. 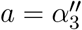; from each pair of the previous step one steady state collide with a third steady state and they disappear. So three saddle-node events happen.
  f. a = α _4_; The triple transcritical event happen.
  g. *a* = *α*_5_; Three saddle-node events happen.
7. 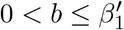; In this case there are five bifurcation points.
  a. *a* = *α*_1_; Three saddle-node events happen.
  b. a = α _2_; Three pitch-fork events happen.
  c. a = α _3_; Three saddle-node events happen.
  d. a = α _4_; The triple transcritical event happen.
  e. a = α _5_; Three saddle-node events happen.

Figures 6a, 6b, 6c, 7a, 7b, and 7c show the behavior of the system for the items 1, 2, 3, 4, (5 and 6), and 7 of the list above respectively. To see the differences between items 5 and 6, see Figures 8 and 9. The differences of these two items are in the steps (5c-5d) and (6c-6e). There are three branches of steady states, each consisting of 3 steady states that are under effect of these bifurcation events. The three steady states of each branch are named by *S*_1_, *S*_2_ and *S*_3_. Among them *S*_2_ is a stable steady state and the rest are unstable. In the left side the steady states are colored with respect to the value of *a* that can be read by the color bar next to it. In the right side there is a schematic plot showing when and which steady states collide and when the two complex conjugate versions of *S*_1_ and *S*_2_ meet and become real again. The horizontal axis shows the value of *a*.

**Figure 8.**
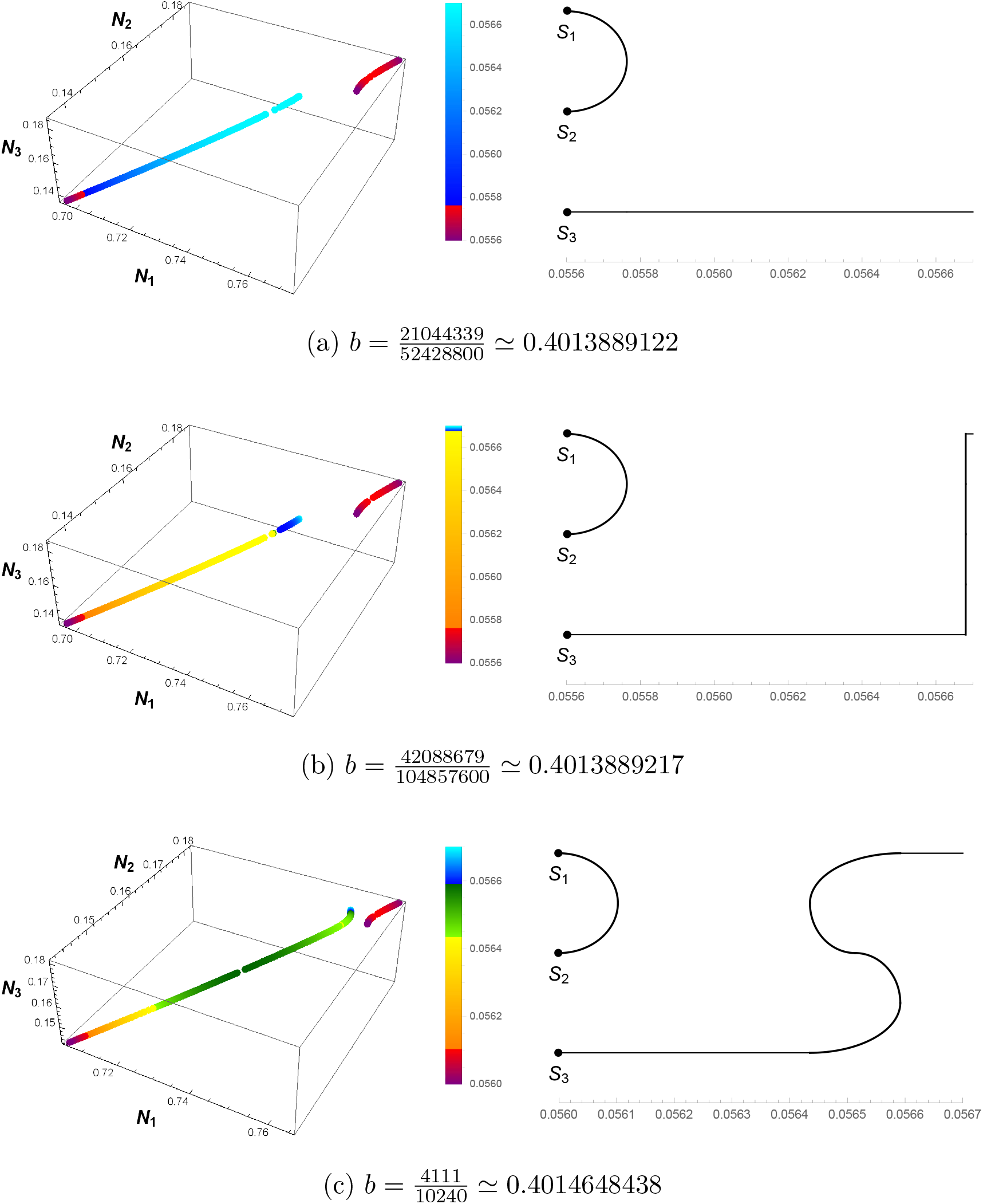
The bifurcation events happening in steps (5c-5d) and (6c-6e) in Section 5. In the left side the sequences of the three involved steady states are depicted for different values of *a*, but a fixed value of *b*. The steady states are colored with respect to their corresponding value of *a* that can be read from the color bars next to the plots. In the right side schematic plots emphasizing when and which steady states collide or return from the complex space to the real space are shown. The value of *a* can be read from the horizontal axis. Continued at Figure 9.

**Figure 9.**
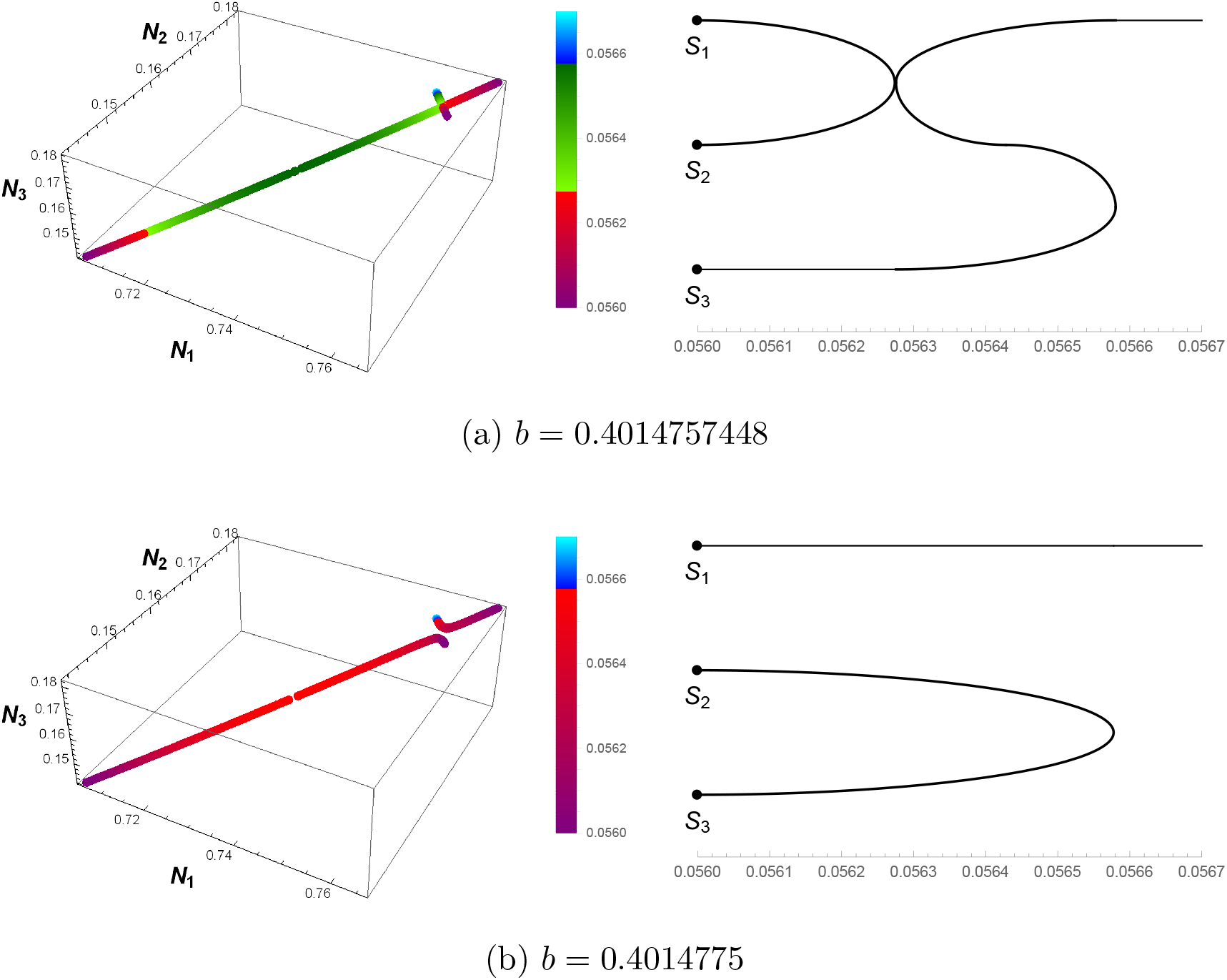
Continuation of Figure 8. The bifurcation events happening in steps (5c-5d) and (6c-6e) in Section 5. In the left side the sequences of the three involved steady states are depicted for different values of *a*, but a fixed value of *b*. The steady states are colored with respect to their corresponding value of *a* that can be read from the color bars next to the plots. In the right side schematic plots emphasizing when and which steady states collide or return from the complex space to the real space are shown. The value of *a* can be read from the horizontal axis.

In Figure 5 the pitchfork bifurcation from Figure 4c is considered. To check if it is a supercritical or subcritical pitchfork bifurcation we computed the eigenvectors of the Jacobian matrix of the system at each steady state and the stability of the steady state along direction of these eigenvectors. The steady states have the same stability behavior along two directions near to the bifurcation point, but different along the third direction. Considering the third direction, the behavior of this pitchfork bifurcation is the same as a supercritical pitchfork. In a supercritical pitchfork bifurcation a stable steady state becomes unstable after the bifurcation event, but there are two stable steady states around it which is termed as “a soft loss of stability” [10, Section 2.7]. Here the event is happening in the opposite direction.

## 6 Conclusion

Allee effect is an important ecological phenomenon, with interesting dynamical properties. To better understand the interplay of Allee effect and population dispersal, we investigated the steady state structure of a system representing three connected populations subject to Allee effect.

To this aim, we used CAD with respect to the discriminant variety, a tool from computational algebraic geometry to find bifurcation points and count the number of steady states. Despite being a very useful tool in theory, its complexity makes it difficult to be used in practice on a normal computer. Here we presented a new algorithm, Algorithm 1 that enabled us to decompose the 2-dimensional parameter space of our model with respect to the number of steady states using 1-dimensional CADs and a search for maximum and minimum of the distances between critical values. We expect that this new approach can be helpful to solve more questions of the same nature in biological context where the exact methods fail due to the computational limits of the computers.

Our results show that even this relatively simple system with three patches can show a great variety of bifurcations. We observed sequences of pitchfork and saddle-node bifurcations, triple-transcritical bifurcations and also a sun-ray shaped bifurcation where twelve steady states meet at a single point then disappear. By increasing the dispersal rate from very small to very large values, one expects that number of steady states will decrease from 27 (completely disconnected patches) to 3 (homogeneous well mixed population), and indeed that happens. However, depending on the Allee threshold, there are sequences of bifurcations for this transition. Interestingly, the number of steady states is not necessarily monotone decreasing from 27 to 3 as we increase dispersal and strengthen the mixing of the populations, but temporarily the number of steady states can go up, as we can find many examples of that in Table 1. Thus, there are situations when increasing the mobility may increase the complexity of the system.

These different pathways are determined by the parameter *b*, which is the ratio of the Allee threshold and the carrying capacity of the populations. From Figure 3 we can observe that for smaller Allee threshold, i.e. when it is easier for a population to establish and survive, a smaller dispersal is sufficient for the system to have only three steady states. By increasing the Allee threshold, the more complex steady state structures exist for a larger range of dispersal parameter, and the patches need to be much more strongly connected to have the simple situation of only three steady states.

The system (2) we studied is a very special case of spatial dispersal models, since we considered identical patches and symmetric spatial movements, to reduce the number of model parameters. For this reason, some of the bifurcations we found are degenerate, and can not be expected to appear in more realistic, noisy population dynamics. We propose as future work to study which bifurcations persist under perturbations of our model. Understanding the spatial patterns of natural populations is a major challenge in ecology, and our results reveal that the interplay of Allee effect and dispersal can create very complex and rich dynamics in this regard.

## Acknowledgements

Gergely Röst was supported by Hungarian grants EFOP-3.6.1-16-2016-00008, NKFIH FK 124016, and TUDFO/47138-1/2019-ITM7. AmirHosein Sadeghimanesh was funded by NKFIH KKP 129877. We thank Matthew England and Tereso Del Río for comments on previous versions of this manuscript.

## Footnotes

1 Computations performed on Windows 10, Intel(R) Core(TM) i7-2670QM CPU @ 2.20GHz 2.20 GHz, x64-based processor, 6.00GB (RAM)

## References

[1] Russell Bradford, James H. Davenport, Matthew England, Hassan Errami, Vladimir Gerdt, Dima Grigoriev, Charles Hoyt, Marek Košta, Ovidiu Radulescu, Thomas Sturm, and Andreas Weber. A Case Study on the Parametric Occurrence of Multiple Steady States. In Proceedings of the 2017 ACM on International Symposium on Symbolic and Algebraic Computation, ISSAC ’17, page 45–52, New York, NY, USA, 2017. Association for Computing Machinery.

[2] Franck Courchamp, Ludek Berec, and Joanna Gascoigne. Allee effects in ecology and conservation. Oxford University Press, 2008.

[3] Wolfram Decker, Gert-Martin Greuel, Gerhard Pfister, and Hans Schönemann. Singular 4-2-0 – A computer algebra system for polynomial computations, 2020. http://www.singular.uni-kl.de.

[4] Matthew England, Hassan Errami, Dima Grigoriev, Ovidiu Radulescu, Thomas Sturm, and Andreas Weber. Symbolic Versus Numerical Computation and Visualization of Parameter Regions for Multistationarity of Biological Networks. In Computer Algebra in Scientific Computing, pages 93–108, Cham, 2017. Springer International Publishing.

[5] Daniel Franco and Alfonso Ruiz-Herrera. To connect or not to connect isolated patches. Journal of Theoretical Biology, 370:72–80, 2015.

[6] Jürgen Gerhard, David J. Jeffrey, and Guillaume Moroz. A package for solving parametric polynomial systems. ACM Communications in Computer Algebra - Sigsam, 43(3/4):61–72, 2010.

[7] Mats Gyllenberg, Jarmo Hemminki, and Toomas Tammaru. Allee effects can both conserve and create spatial heterogeneity in population densities. Theoretical population biology, 56(3):231–242, 1999.

[8] Heather A. Harrington, Dhagash Mehta, Helen M. Byrne, and Jonathan D. Hauenstein. Decomposing the Parameter Space of Biological Networks via a Numerical Discriminant Approach. Maple in Mathematics Education and Research. MC 2019. Communications in Computer and Information Science, 1125:114–131, 2020.

[9] Alan Hastings and Frank M. Hilker. Multiple attractors and long transients in spatially structured populations with an Allee effect. Bulletin of Mathematical Biology, 82(6):1522–9602, 2020.

[10] John K. Hunter. Introduction to Dynamical Systems. https://www.math.ucdavis.edu/~hunter/m207/m207.pdf, 2011.

[11] Diána Knipl and Gergely Röst. Spatially heterogeneous populations with mixed negative and positive local density dependence. Theoretical Population Biology, 109:6–15, 2016.

[12] Daniel Lazard and Fabrice Rouillier. Solving Parametric Polynomial Systems. Journal of Symbolic Computation, 42(6):636–667, 2007.

[13] Guillaume Moroz. Sur la décomposition réelle et algébrique des systémes dépendant de paramétres. PhD thesis, Université Pierre et Marie Curie - Paris VI, 2008. https://tel.archives-ouvertes.fr/tel-00812436/file/these_moroz.pdf.

[14] Mariano Sigman and Gabriel B. Mindlin. Dynamics of Three Coupled Excitable Cells with D3 Symmetry. International Journal of Bifurcation and Chaos, 10(7):1709–1728, 2000.

